# Discovery of Novel Glycerolated Quinazolinones from *Streptomyces* sp. MBT27

**DOI:** 10.1101/484535

**Authors:** Nataliia Machushynets, Changsheng Wu, Somayah S. Elsayed, Thomas Hankemeier, Gilles P. van Wezel

**Affiliations:** Molecular Biotechnology, Institute of Biology, Leiden University, Sylviusweg 72, 2333 BE, The Netherlands; State Key Laboratory of Microbial Technology, Institute of Microbial Technology, Shandong University, Qingdao 266237, P.R. China; Leiden Academic Centre for Drug Research (LACDR), Leiden University, Einsteinweg 55, 2333 CC, Leiden, The Netherlands

**Keywords:** 4(3*H*)-quinazolinones, molecular networking, metabolomics, carbon source

## Abstract

Actinobacteria are a major source of novel bioactive natural products. A challenge in the screening of these microorganisms lies in finding the favorable growth conditions for secondary metabolite production and dereplication of known molecules. Here, we report that *Streptomyces* sp. MBT27 produces 4-quinazolinone alkaloids in response to elevated levels of glycerol, whereby quinazolinones A (**1**) and B (**2**) form a new sub-class of this interesting family of natural products. Global Natural Product Social molecular networking (GNPS) resulted in a quinazolinone-related network that included anthranilic acid (**3**), anthranilamide (**4**), 4(3H)-quinazolinone (**5**) and 2,2-dimethyl-1,2-dihydroquinazolin-4(3H)-one (**6**). Actinomycins D (**7**) and X2 (**8**) were also identified in the extracts of *Streptomyces* sp. MBT27. The induction of quinazolinone production by glycerol combined with biosynthetic insights provide evidence that glycerol is integrated into the chemical scaffold. The unprecedented 1,4-dioxepane ring that is spiro-fused into the quinazolinone backbone, is most likely formed by intermolecular etherification of two units of glycerol. Our work underlines the importance of varying the growth conditions for the discovery of novel natural products and for understanding their biosynthesis.

## Introduction

Actinobacteria are a major source of bioactive compounds, producing some two thirds of all antibiotics as well as molecules with a wide variety of activities such as anticancer, antifungal and immunosuppressant [2,4]. Traditionally, microbial natural product (NP) discovery has been done via high-throughput screening followed by iterative bioassay-guided fractionation and structure elucidation. While such pipelines were extremely successful and delivered a plethora of therapeutic agents, in the modern era the large pharmaceutical companies moved out of NP-discovery programs due to high cost and chemical redundancy [1,9,29]. At the same time, the power of genome sequencing brought the recognition that microorganisms harbor a vast and yet untapped biosynthetic potential, and it rapidly became clear that the potential for metabolic diversity of even the best-studied model organisms as producers of natural products had been grossly underestimated [3,11,19]. How could these compounds have been missed by the very extensive HT screening campaigns of the 20th century? The answer is that many of the biosynthetic gene clusters (BGCs) discovered by genome mining are poorly expressed or cryptic under laboratory conditions [22,28]. A drug-discovery pipeline that is rapidly gaining momentum involves combining genome mining with fluctuating the culturing conditions to achieve differential synthesis of NPs, followed by the metabolic profiling-based identification of the bioactivity of interest [13,16,48,50]. A major challenge thereby lies in finding the appropriate chemical triggers or ecological cues to elicit the production of cryptic antibiotics (recently reviewed in [34,55,56]). The use of chemical elicitors is thereby a promising approach [10,42,56].

Manipulation of fermentation conditions for promising producer strains, known as the “one strain many compounds” (OSMAC) approach, is an effective way of enhancing the production of secondary metabolites [33,5]. The regulatory networks that control the production of bacterial natural products respond strongly to changes in carbon, nitrogen or phosphate concentration [41,40]. The reisolation of known metabolites is a major bottleneck in the discovery of new bioactive natural products. A crucial step in this regard is the early identification of already known substances, in order to concentrate the efforts on the discovery of new ones, a process known as dereplication [15]. Current dereplication strategies include hyphenated techniques, such as LC-MS, LC-NMR, LC-NMR-MS, and LC-SPE-NMR. Bioactivity fingerprinting has also been used to dereplicate natural products based on their biological modes of action [36], while molecular networking is a powerful tool for the visualization and dereplication of natural products [54,46]. Mass spectrometry-based molecular networking relies on clustering of molecules based on similarities in their MS/MS fragmentation patterns, which depends on the structural features of the ionized molecules. The resulting clusters allow scientists to visually explore the metabolites produced by a given strain under a specific growth condition, allowing rapid dereplication of known compounds by automated spectral library searches, and to visualize their unknown structural analogues [45].

In this work we analyzed the potential of *Streptomyces* sp. MBT27 as a producer of natural products in response to changes in the carbon source. The strain had previously been identified as a promising producer of NPs [57]. Extensive fluctuations in the secondary metabolite profiles were observed depending on the carbon source used, and statistical methods combined with GNPS molecular networking identified a family of known as well as novel quinazolinone compounds, in response to high concentrations of glycerol. Quinazolinones are heterocyclic compounds with a wide range of medical applications, such as antimicrobial, antiviral, antituberculosis and as enzyme inhibitors (reviewed in [24,17,20]). Combination of MS and NMR methods identified the novel quinazolinones A (1) and B (2), which further expands the chemical space of this rich family of natural products.

## Materials and Methods

### Bacterial strains and growth conditions

*Streptomyces* sp. MBT27 was obtained from the Leiden University strain collection and had previously been isolated from the Qingling Mountains, Shanxi province, China [57]. Cultures were grown in triplicate in 100 mL Erlenmeyer flasks with 30 mL of liquid minimal medium (MM; [21]), supplemented with various carbon sources, and inoculated with 10 μL of 10^9^/mL spore suspension. The carbon sources (percentages in w/v) were: 1% mannitol + 1% glycerol, 1% mannitol, 2% mannitol, 1% glycerol, 2% glycerol, 1% glucose, 2% glucose, 1% fructose, 1% arabinose or 1% N-acetylglucosamine (GlcNAc). The cultures were incubated in a rotary shaker at 30 °C at 220 rpm for seven days.

### General Experimental Procedures

NMR spectra were recorded in deuterated methanol (CD_3_OD) on a Bruker 600 MHz, and were referenced using the residual ^1^H signal of deuterated solvent at 3.30 ppm [49,53]. FT-IR was measured on Perkin-Elmer FT-IR Spectrometer Paragon 1000. UV measurements were performed using a Shimadzu UV mini-1240. Optical rotations were measured on a JASCO P-1010 polarimeter. HPLC purification was performed on Waters preparative HPLC system comprised of 1525 pump, 2707 autosampler, 2998 PDA detector, and Water fraction collector III. The columns used were SunFire C_18_ column (5 μm, 100 Å, 10 × 250 mm) and SunFire C_18_ column (10 μm, 100 Å, 19 × 150 mm). TLC was performed using aluminum plates coated with silica gel 60 F_254_ (Merck). All organic solvents and chemicals were of analytical or LCMS grade, depending on the experiment.

### Metabolite profiling

Following fermentation, culture supernatants were extracted with ethyl acetate (EtOAc) and evaporated under reduced pressure. For LC-ESI-QTOFMS analyses, extracts were dissolved in MeOH to a final concentration of 1 mg/mL, and 1 μL was injected into Waters Acquity UPLC system equipped with Waters Acquity HSS C_18_ column (1.8 μm, 100 Å, 2.1 × 100 mm), which is coupled to Agilent 6530 QTOF MS equipped with Agilent Jet Stream ESI source (Agilent Technologies, Inc., Palo Alto, CA, USA). For the LC, solvent A was 95% H_2_O, 5% acetonitrile (ACN) and 0.1% formic acid; solvent B was 100% ACN and 0.1% formic acid. The gradient used was 2% B for 1 min, 2–85% for 9 min, 85–100% for 1 min, and 100% for 3 min. The flow rate used was 0.5 mL/min. As for the MS, the following ESI source parameters were used: capillary voltage 3 kV, source temperature 325 °C, drying gas flow rate 10 L/min, and fragmentor 175 V. Full MS spectra were acquired in positive mode in the range of 100-1700 *m/z,* in the extended dynamic range mode. Internal reference masses of purine and Agilent HP-921 were continuously delivered to the ESI source through an Agilent 1260 isocratic pump.

Thermo Instruments MS system (LTQ Orbitrap XL, Bremen, Germany) equipped with an electrospray ionization source (ESI) was used for LC-MS/MS analysis. The Waters Acquity UPLC system equipped with Waters Acquity PDA was run using a SunFire Waters C_18_ column (3.5 μm, 100 Å, 4.6 × 150 mm), at a flow rate of 0.9 mL/min. Solvent A was 95% H_2_O, 5% acetonitrile (ACN) and 0.1% formic acid; solvent B was 100% ACN and 0.1% formic acid. The gradient used was 2% B for 1 min, 2-85% for 15 min, 85–100% for 3 min, and 100% for 3 min. As for the MS, the following ESI parameters were used: capillary voltage 5 V, spray voltage 3.5 kV, capillary temperature 300 °C, auxiliary gas flow rate 10 arbitrary units, and sheath gas flow rate 50 arbitrary units. Full MS spectra were acquired in the Orbitrap in positive mode at a mass range of 100–2000 *m/z,* and FT resolution of 30000. Data dependent MS^2^ spectra were acquired in the ion trap for the three most intense ions using collision induced dissociation (CID). The resulting chemical data were compared with those in SciFinder and Antibase [25].

### Large-scale fermentation and isolation of metabolites (1) and (2)

For large-scale fermentation, *Streptomyces* sp. MBT27 was grown in eight 2 L Erlenmeyer flasks, each containing 500 mL liquid MM supplemented with 2% w/v glycerol at 30 °C for seven days. The metabolites were extracted from the spent media of culture filtrates using EtOAc, and the solvent was subsequently evaporated under reduced pressure at 40^0^C. The crude extract (1.4 g) was adsorbed onto 1.4 g silica gel (pore size 60 Å, 70-230 mesh, Sigma Aldrich), and loaded on silica column, which was eluted using gradient mixtures of *n*-hexane, acetone, and MeOH. The fractions eluted with acetone 100% were combined, reconstituted in MeOH, and injected into the preparative SunFire column (19 x 150 mm), which was eluted with H_2_O:MeOH gradient of 7–100% in 30 min, at a flow rate of 15 mL/min, to yield five fractions. Quinazolinones A and B were further purified on the semipreparative SunFire column (10 x 250 mm), run at 3 mL/min. Fraction 2 was eluted using H_2_O:MeOH gradient of 20–40% in 20 min, to yield Quinazolinone A (**1, 1** mg). On the other hand, Fraction 3 was eluted using H_2_O:MeOH gradient of 30-40% in 20 min, to yield Quinazolinone B (2, 1.5 mg).

#### Quinazolinone A (1)

colorless, amorphous powder; [α]_D_^20^ 2.3 (c 0.1, MeOH); UV (MeOH) λ_max_ (log ε) 224 (4.09), 322 (2.98) nm; IR V_max_ 3334, 2922, 1652, 1475, 1052 cm^-1^; ^1^H and ^13^C NMR data, see Table 1; HRESIMS (positive mode) *m/z* 281.1130 [M + H]+ (calcd. for C_13_H_17_N_2_O_5_, 281.1132).

**Table 1.** ^1^H and ^13^C NMR data for compounds 1 and 2^*a*^

| NO. | **1** |  | **2** |  |
| --- | --- | --- | --- | --- |
| | $\delta_C$ | $\delta_H$ (mult., $J$ in Hz) | $\delta_C$ | $\delta_H$ (mult., $J$ in Hz) |
| 2 | 79.0 |  | 73.3 |  |
| 4 | 166.1 |  | 166.7 |  |
| 4a | 121.8 |  | 114.5 |  |
| 5 | 128.8 | 7.88 (dd, $J = 7.8, 1.2$) | 128.5 | 7.66 (dd, $J = 7.8, 1.2$) |
| 6 | 123.0 | 7.07 (t, $J = 7.8$) | 118.5 | 6.69 (td, $J = 7.8, 1.2$) |
| 7 | 134.6 | 7.46 (td, $J = 7.8, 1.2$) | 135.3 | 7.26 (td, $J = 7.8, 1.2$) |
| 8 | 122.3 | 7.23 (brd, $J = 7.8$) | 115.8 | 6.73 (brd, $J = 7.8$) |
| 8a | 143.7 |  | 148.6 |  |
| 1' | 73.9 | 4.43 (d, $J = 9.0$); 4.24 (d, $J = 9.0$) | 64.5 | 3.67 (d, $J = 10.8$); 3.62 (d, $J = 10.8$) |
| 2' | 64.6 | 3.59 (d, $J = 11.4$); 3.47 (d, $J = 11.4$) | 64.5 | 3.67 (d, $J = 10.8$); 3.62 (d, $J = 10.8$) |
| 1'' | 65.7 | 3.86 (d, $J = 12.6$); 3.80 (d, $J = 12.6$) | | |
| 2'' | 102.9 |  |  |  |
| 3'' | 61.9 | 3.66 (d, $J = 3.0$) | | |

<sup>a</sup> **1** and **2** were recorded in CD<sub>3</sub>OD, at 298 K. All chemical shift assignments were done on the basis of 1D and 2D NMR techniques.

#### Quinazolinone B (2)

colorless, amorphous powder, UV (MeOH) λ_max_ (log ε) 226 (4.18), 347 (3.03) nm; IR V_max_ 3350, 2950, 1649, 1526, 1049, 751 cm^-1^; ^1^H and ^13^C NMR data, see Table 1; HRESIMS (positive mode) *m/z* 209.0915 [M + H]+ (calcd. for C_10_H_13_N_2_O_3_, 209.0921).

### Computation of mass spectral networks

MS/MS raw data were converted to a 32 bit mzXML file using MSConvert (ProteoWizard) [6] and spectral networks were assembled using Global Natural Product Social molecular networking (GNPS) (https://gnps.ucsd.edu) as described [45]. For both parent and MS/MS fragment ions, the mass tolerance was set to 0.5 Da, while the minimum cosine score was set to 0.7. The data were clustered using MSCluster with a minimum cluster size of three spectra. The spectra in the network were also searched against GNPS spectral libraries. A minimum score of 0.5 was set for spectral library search, with at least two fragment peaks matching. Cytoscape 3.5.1. was used for visualization of the generated molecular networks [38]. The edge thickness was set to represent the cosine score, with thicker lines indicating higher similarity between nodes. LC-MS/MS data were deposited in the MassIVE Public GNPS data set (MSV000082988). The molecular networking job in GNPS can be found at https://gnps.ucsd.edu/ProteoSAFe/status.jsp?task=8fd1fcfa0ff744a9808e80bc7be12115. The annotated MS/MS spectra were deposited in the GNPS spectral library for quinazolinone A (<u>CCMSLIB00004684355</u>), and B (<u>CCMSLIB00004684354</u>).

### Statistical analysis

Prior to statistical analysis, mzXML files were imported into Mzmine 2.31 [30] for data processing. Mass ion peaks were detected using the exact mass algorithm with a noise level set to 1.0 × 10^4^. Afterwards, chromatograms were built for the detected masses with a minimum time span of 0.05 min, *m/z* tolerance of 0.001 *m/z* and minimum height of 1.0 × 10^4^. Chromatogram deconvolution was then performed using local minimum search algorithm (search minimum in RT range 0.1 min, chromatographic threshold 90%, minimum relative height 1%, minimum absolute height 1.0 × 10^4^, minimum ratio of peak top/edge 2 and peak duration range 0.05–3 min). In the generated peak lists, isotopes were identified using isotopic peaks grouper (m/z tolerance 0.001 *m/z* and retention time tolerance 0.1 min), and variations in retention time were reduced using retention time normalizer (m/z tolerance 0.001 *m/z* and retention time tolerance 1 min). All the peak lists were subsequently aligned using join aligner (m/z tolerance 0.001 m/z, *m/z* weight 20, retention time tolerance 0.1 min, and retention time weight 20), and missing peaks were detected through gap filling using peak finder (intensity tolerance 1.0%, *m/z* tolerance 0.001 m/z, and retention time tolerance of 0.2 min). Finally, the aligned peak list was exported as a comma separated file for statistical analysis.

Statistical analysis was performed using MetaboAnalyst [7], where log transformation and pareto scaling was initially applied to the data. A heat map of all detected masses, among the different growth conditions, was generated in MetaboAnalyst, to which additional hierarchical clustering analysis (HCA) was performed using Euclidean distance measure and Ward clustering algorithm. Student’s t-tests with multiple testing correction (Benjamini Hochberg false discovery rate or FDR) were used to determine significant differences in the intensities of the metabolites, under two different growth conditions. The thresholds set for statistically significant differences were a fold change ≥ 4, together with FDR corrected *p*-value ≤ 0.05 Based on these criteria, a volcano plot was generated. To identify the difference in intensity of a single mass feature among multiple growth conditions, one-way ANOVA was performed, followed by a *post hoc* Tukey’s honest significant difference (HSD) test.

## RESULTS AND DISCUSSION

### The influence of carbon sources on secondary metabolite production

Previous screening of our in-house actinomycete collection, obtained from remote mountain soils, showed it is a promising source of new bioactive compounds [57]. Under specific growth conditions, these isolates exert potent inhibitory activity against the so-called ESKAPE pathogens [31] *Enterococcus faecium, Staphylococcus aureus, Klebsiella pneumoniae, Acinetobacter baumannii, Pseudomonas aeruginosa* and *Enterobacter* spp. In the current study, the potential of one of the *Streptomyces* strains in our collection, namely *Streptomyces* sp. MBT27, was analyzed to study the effect of carbon sources on its metabolite profile. Traditional approaches such as changes in fermentation conditions, are known to induce significant changes in the microbial metabolome. Culture medium components, and particularly the carbon source, have major effects on the production of secondary metabolites [43,35,5,26]. The differential production of small molecules is ideal for metabolomics studies, whereby metabolic variations are correlated statistically to bioactivity, thus facilitating the identification of the bioactive molecule of interest [48,51,52]. To establish the potential of *Streptomyces* sp. MBT27, the strain was grown in liquid minimal media containing different carbon sources, namely (percentages in w/v): 1% of both mannitol and glycerol, 1% mannitol, 2% mannitol, 1% glycerol, 2% glycerol, 1% glucose, 2% glucose, 1% fructose, 1% arabinose, or 1% N-acetylglucosamine (GlcNAc). The latter is an elicitor of antibiotic production, via metabolic interference with the global nutrient sensory network controlled by DasR [39,32]. Phosphate buffer was omitted from the MM as it repressed the production of secondary metabolites by *Streptomyces* sp. MBT27 (data not shown). To identify the secondary metabolites in the cultures, supernatants were extracted with EtOAc, and the resulting crude extracts subjected to LC-MS analysis. Using the LC-MS data, a heat map with added hierarchical clustering was generated, to visualize the production of different metabolites under different culture conditions (Fig. 1). Hierarchical clustering analysis of the LC-MS data allows effective comparative analysis of metabolomics data, and the heat map revealed major differences in the metabolic profiles of *Streptomyces* sp. MBT27, whereby different groups of metabolites were enhanced depending on the carbon source used. Interestingly, not only the type of carbon source, but especially also the concentration resulted in large changes in the metabolic profiles. Doubling the concentration of either glycerol or glucose from 1% to 2% had a profound effect on the metabolic profile. Thin layer chromatography (TLC) was conducted to compare metabolic profiles of the 1% and 2% glycerol-grown cultures. Interestingly, this revealed that several fluorescent compounds were differentially produced in the extracts of 2% glycerol-grown cultures relative to those produced in 1% glycerol (Fig. 2a). In order to provide statistical relevance to the data, the metabolic profiles of 1% and 2% glycerol-grown cultures were compared using a volcano plot (Fig. 2b). The volcano plot was then searched for the mass features which increased in production in 2% glycerol as compared to 1%. A mass of *m/z* 281.1151 (1) stood out as its intensity had increased by around 7000 fold (p-value = 0.002) in cultures fermented in 2% glycerol as compared to 1%.

**Fig 1.**
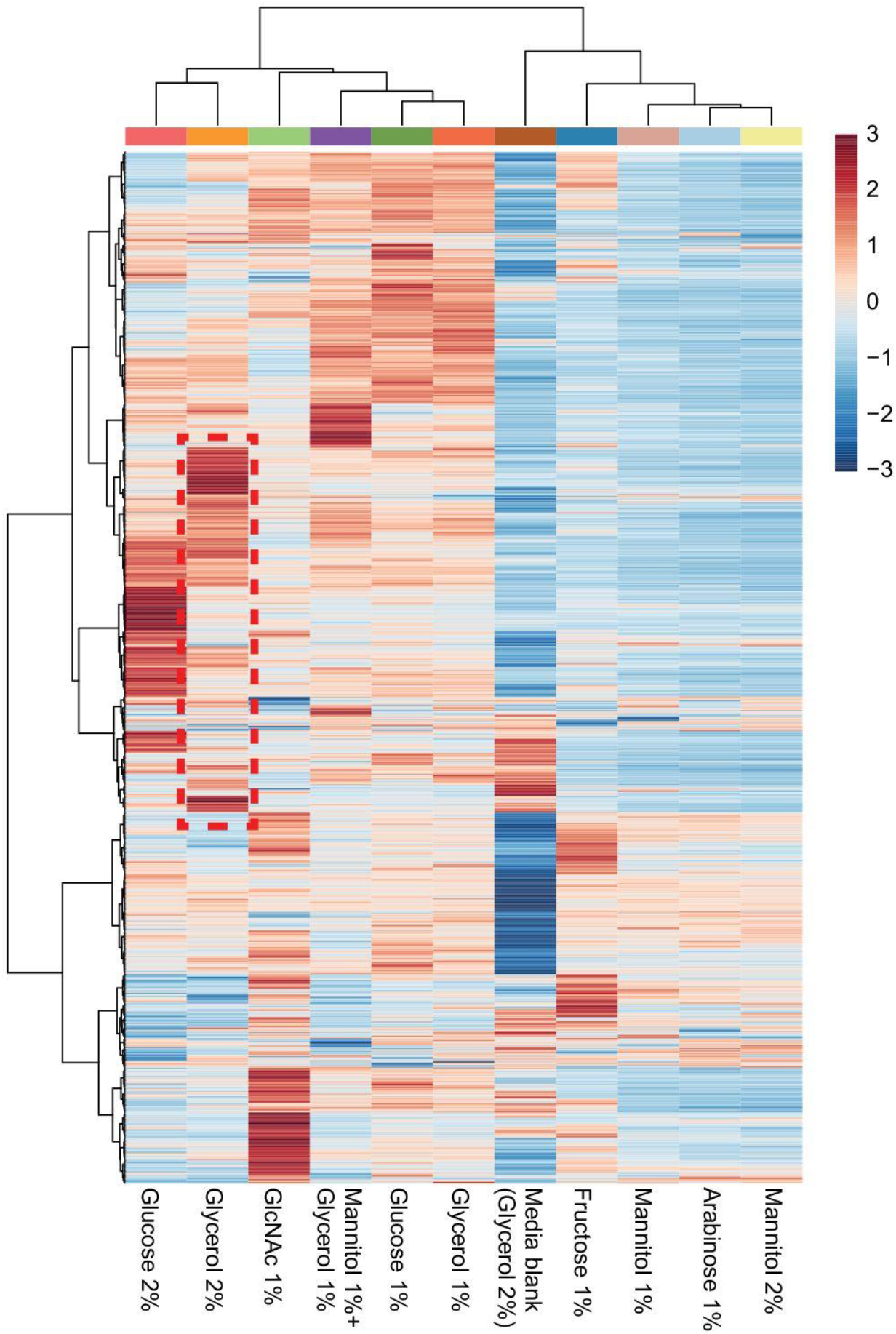
Analysis of secondary metabolites produced by *Streptomyces* sp. MBT27. The heat map depicts the relative abundance of the metabolites (rows) produced under different growth conditions (columns)

**Fig 2.**
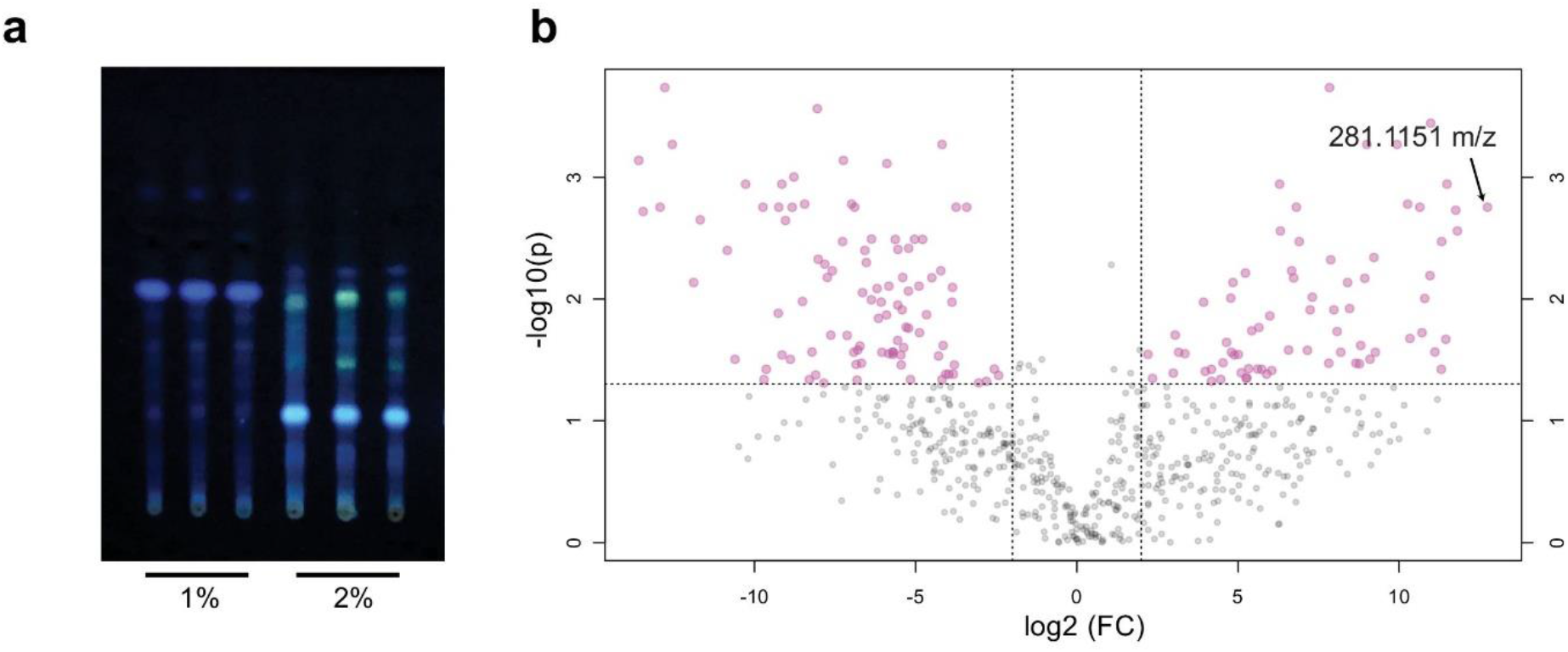
Secondary metabolite profiles found in 1% and 2% glycerol-grown cultures of *Streptomyces* sp. MBT27. **a** Thin layer chromatography of extracts from the cultures grown in MM with either 1% or 2% glycerol as the carbon source. **b** Volcano plot highlighting the differences in metabolite profiles. The *x* and *y* axes of the volcano plot represent the log2 fold changes and the corresponding −log10 FDR-adjusted *p*-value of all metabolites, respectively. Pink circles represent metabolites with an intensity difference of more than 4-fold (*p*-value ≤ 0.05). Ions present in the left and right quadrants are associated with the 1% and 2% glycerol-grown cultures, respectively. Metabolites situated towards the left and right top quadrants represent values of large magnitude fold changes as well as high statistical significance.

Global Natural Product Social (GNPS) molecular networking [45] was employed to detect the MS/MS structural relatedness among molecules in an automated manner; the software generates a molecular network wherein molecules with related scaffolds cluster together [45]. A network representing the ions detected in the crude extract of *Streptomyces sp.* MBT27 grown with 2% glycerol was constructed, revealing 183 nodes clustered in 19 spectral families (Fig. 3). GNPS dereplication based on matching with its MS/MS spectral database highlighted some known metabolites. These included anthranilic acid (**3**), anthranilamide (**4**) [37], actinomycin D (**7**), and actinomycin X2 (**8**) [44]. The annotation of the compounds was supported by comparison of the exact mass, fragmentation pattern, and UV spectra, with reference data. Moreover, crude extracts of MBT27 possessed antimicrobial activity against *B. subtilis*, which was most likely due to the expression of actinomycins (data not shown). Additional metabolites could be dereplicated through manual comparison of their spectral data against the microbial natural products database Antibase. These previously described metabolites were 4(3*H*)-quinazolinone (**5**) [23] and 2,2-dimethyl-1,2-dihydroquinazolin-4(3*H*)-one (**6**) [8].

**Fig 3.**
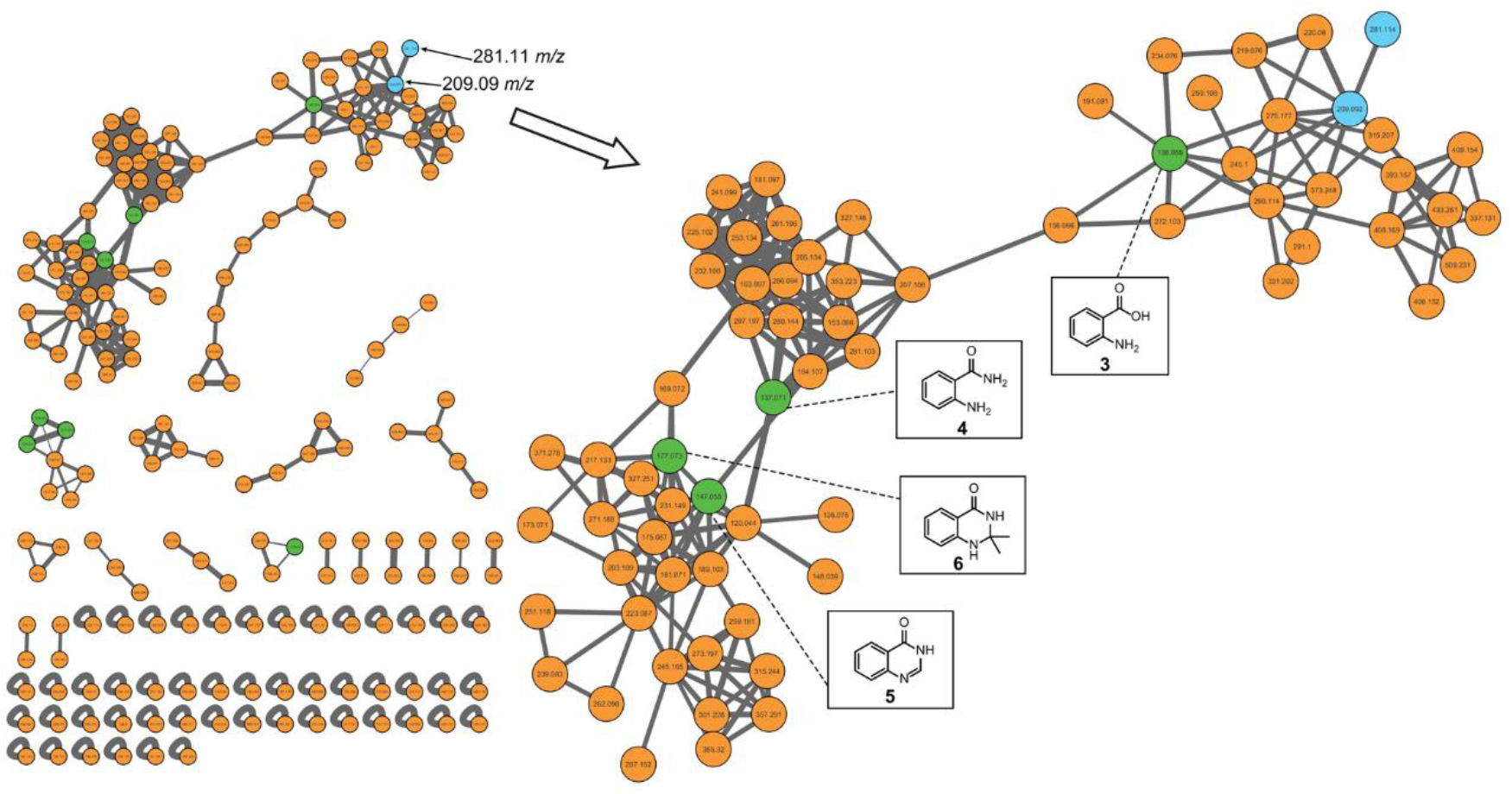
GNPS molecular network of the ions detected in the crude extract of *Streptomyces* sp. MBT27 grown in MM with 2% glycerol. Enlargment is made to the cluster including new metabolites. Orange nodes represent all the ions detected in the extract. Green nodes represent the dereplicated metabolites, while blue nodes represent the ions upregulated when glycerol concentration was increased from 1% to 2%.

Besides known molecules, the network also contained many mass features that could not be assigned to any of the previously identified metabolites. One of these was an ion with an *m/z* value of 281.1151 (1), which was very highly increased in intensity (7000 fold, *p*-value = 0.002) when the glycerol concentration was increased from 1% to 2%. It was closely connected (cosine score 0.91) to another upregulated ion with an *m/z* value of 209.0932 (**2**), which was also not previously reported. Both ions were part of the large spectral family comprising the annotated anthranilic acid (**3**), anthranilamide (**4**), and the 4-quinazolinones **5** and **6,** suggesting that they are structurally related metabolites. It is important to note that the concentrations of quinazolinones A and B did not change significantly when the concentration of either mannitol or glucose was doubled from 1% to 2%, or when mannitol was added as additional carbon source to glycerol (Fig. S1). This strongly suggests that the effect of glycerol is not due to increased availability of the carbon source.

The relationship between the increase in glycerol concentration and the strong increase in the production of the compounds **1** and **2**, was investigated further by expanding the range of glycerol concentrations and analyzing the metabolic profiles. For this, *Streptomyces* sp. MBT27 was cultured in glycerol concentrations ranging from 1-4% (w/v). ANOVA, followed by a *post hoc* Tukey’s HSD test, was performed to trace the variation in the production of **1** and **2** among the different culturing conditions. A box plot was used to visualize such variation (Fig. S2). As observed earlier, the production of **1** and **2** was significantly increased when the glycerol concentration was increased from 1% to 2%. However, further increase in glycerol concentration (3% and 4%), did not lead to any significant increase in the production of **1** and **2**. Accordingly, MM with 2% glycerol was used to culture the bacteria, for the purpose of purification and identification of the new metabolites **1** and **2**.

### Isolation and structure elucidation of novel quinazolinones

To elucidate the structure of **1** and **2**, large-scale fermentation was done to obtain larger quantities of the compounds. For this, *Streptomyces* sp. MBT27 was fermented in a total of 4 L of liquid MM supplemented with 2% glycerol, and the supernatant was extracted with EtOAc. Following repeated chromatographic isolation, compounds **1** and **2** were obtained as pure, colorless, amorphous powders. Both compounds were fluorescent, showing UV absorption maxima at 224 and 322 nm and at 226 and 347 nm, respectively. The final structures of **1** and **2** were determined by the combination of NMR and high resolution MS (Fig. 4).

**Fig 4.**
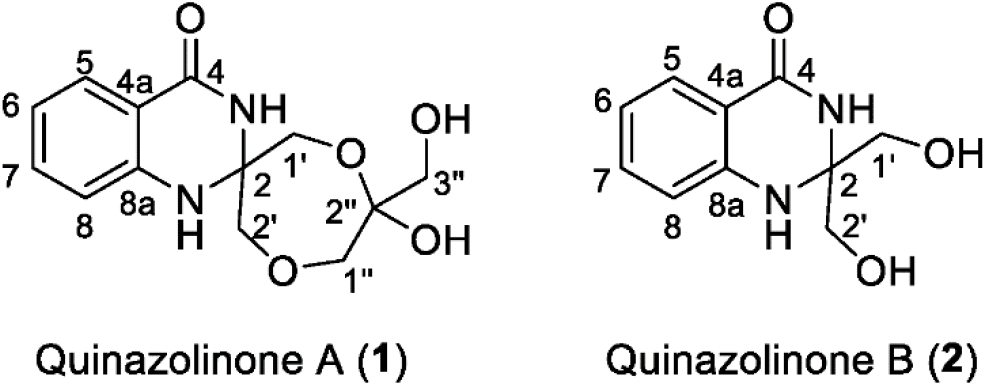
Chemical structures of quinazolinones A (**1**) and B (**2**)

The high resolution mass of *m/z* 281.1151 for an [M + H]+ established a molecular formula of C_13_H_16_N_2_O_5_, with seven degrees of unsaturation, for 1 (yield 0.25 mg/L). The deduced molecular formula was corroborated by the ^13^C NMR attached proton test (APT) spectrum that exhibited 13 carbons in total. On the other hand, the ^1^H NMR spectrum of **1** (Table 1) presented four coupling aromatic signals at *δ_H_* 7.88 (dd, *J* = 7.8, 1.2, H-5), 7.07 (t, *J* = 7.8, H-6), 7.46 (td, *J* = 7.8, 1.2, H-7), and 7.23 (brd, *J* = 7.8, H-8), indicating an o-disubstituted aromatic benzene system. The HMBC correlation (Fig. 5) from H-5 to C-4 (*δ*_c_ 166.1) confirmed one of substituents on the benzene ring to be an ester/amide carbonyl group, while the downfield chemical shift of C-8a at *δ*_c_ 143.7 indicated it was nitrogenated. Four O-bearing methylene groups at SH 4.43 (d, *J* = 9.0, H-1’a) and 4.24 (d, *J* = 9.0, H-1’b); 3.59 (d, *J* = 11.4, H-2’a) and 3.47 (d, *J* = 11.4, H-2’b); 3.86 (d, *J* = 12.6, H-1”a) and 3.80 (d, *J* = 12.6, H-1″b); and 3.66 (d, *J* = 3.0, H2-3″) were resolved by HSQC experiments. Three of these methylene groups were part of a 1,4-dioxepane ring system, which was established based on the HMBC correlations observed from H-1’ to C-2 (*δ*_c_ 79.0) and C-2’ (*δ*_c_ 64.6), from H-2’ to C-1″ (*δ*_c_ 65.7), and from H-1’ to the hemiacetal C-2″ (*δ*_c_ 102.9). The remaining oxymethylene group CH_2_-3″ was connected to C-2″ based on the HMBC correlations observed from H_2_-3″ to C-1″ and C-2″. The two substructures obtained accounted for all of the oxygens and six out of the seven degrees of unsaturation required by the molecular formula of **1**. Accordingly, an additional ring including two nitrogen atoms was deduced to connect the two substructures, forming a 4-quinazolinone ring. Consequently, compound 1 was identified to be a *di*-glycerolated 4-quinozolinone, and was named quinazolinone A.

**Fig 5.**
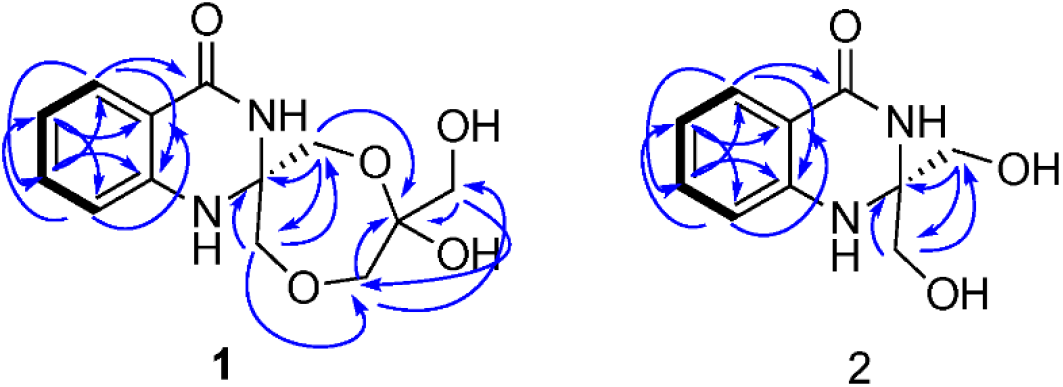
Key COSY (━) and HMBC (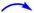) correlations for 1 and 2

The [M + H]+ molecular ion at *m/z* 209.0932 in the ESI-HRMS spectrum resulted in a molecular formula of C_10_H_12_N_2_O_3_ for **2** (yield 0.3 mg/L). The ^13^C NMR spectrum presented 10 signals, which was consistent with the molecular formula (Table 1). The ^1^H NMR spectrum of 2 resembled that of **1** in the downfield aromatic region, and the major difference is the absence of a set of signals for the additional glycerol unit. Further HMBC experiment confirmed 2 is a mono-glycerolated 4-quinazolinone (Fig. 5). Compound **2** was named quinazolinone B.

The 4-quinazoline alkaloids represent an important class of nitrogen-containing heterocyclic compounds [27], which have highly diverse biological activities, such as antitumor, anti-inflammatory, antihypertensive, antimicrobial, anticonvulsant and antifungal activities [18]. Though the basic bicyclic core (the fused benzene ring and pyrimidine ring) is normally conserved in most of naturally occurring 4-quinazoline derivatives [27], a variety of structural substitutions have oftentimes been found on the pyrimidine ring of the 4-quinazolinone. The chemical structures of quinazolinones A and B and the glycerol-dependent production suggest that glycerol participates in the construction of the ring system. We propose that quinazolinones A and B form a new sub-branch in the family of the quinazolines. Particularly, quinazolinone A contains an exciting seven-membered ring that we believe may be formed by intermolecular etherification of two units of glycerol, and this ring is further linked with the quinazolinone backbone by a spiro atom at C-2, which is unprecedented. Antimicrobial activities of compounds **1** and **2** were tested against *B. subtilis* and *E. coli.* None of the compounds showed antimicrobial activity against these indictor strains (data not shown).

### Biosynthesis of quinazolinones A and B

In our study, the production of 1 and 2 was dramatically enhanced in the cultures with 2% glycerol. Elicitation by glycerol is an indication that glycerol may play a key role during their biosynthesis. The aromatic ring system of quinazoline alkaloids is known to be derived from anthranilic acid, which in turn is biosynthesized through the shikimate pathway [12]. Detection of anthranilamide in the extract of *Streptomyces* sp. MBT27 leads to the assumption that the biosynthesis of the isolated quinazolinones starts from anthranilic acid, which is then converted to anthranilamide (Fig. 6). Further successive attachments of two molecules of glycerol then results in the formation of **2**, followed by **1**. However, multiple biosynthetic routes have been proposed for quinazolinones, because the C-2 residue of the quinazoline ring may originate from various precursors. For example, the C-2 and the remaining non-aromatic part of the quinazolinone alkaloid chrysogine, produced by *Penicillium chrysogenum,* are biosynthesized from pyruvic acid via an NRPS system [47], while the non-aromatic part of the quinazoline alkaloid peganine, produced by the plant *Peganum harmala,* is derived from ornithine [12]. Furthermore, fumiquinazoline F originates from a fungal nonribosomal peptide synthetase (TqaA) and is biosynthesized from anthranilic acid, L-tryptophan and L-alanine [14].

**Fig 6.**
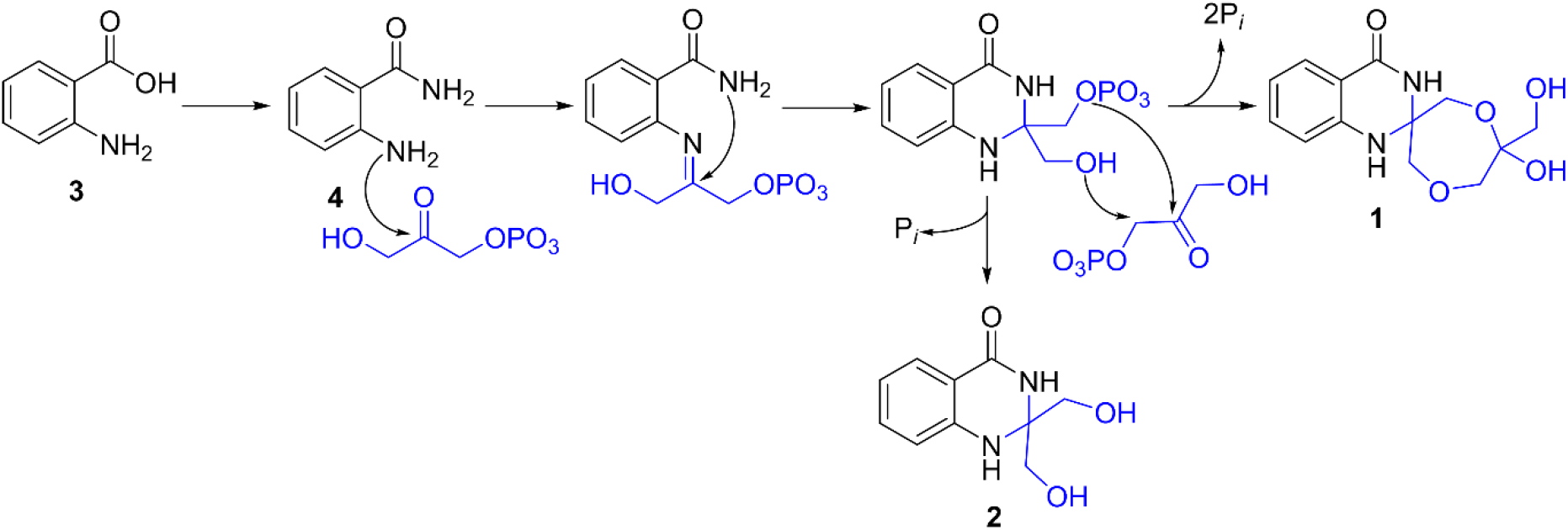
Proposed biosynthetic pathway for quinazolinone A (**1**) and B (**2**)

In conclusion, the metabolic potential of *Streptomyces* sp. MBT27 was analyzed under different nutritional conditions, showing major changes in the metabolome, depending on the carbon source used. In particular, the relatively unspectacular change from 1% to 2% glycerol resulted in a surprisingly global change in the secondary metabolome, with some compounds changing by almost four orders of magnitude. The use of GNPS molecular networking, together with LC-MS based dereplication, allowed us to annotate a cluster of quinazolinone-family molecules that connects to anthranilic acid and anthranilamide. Isolation and structure elucidation revealed the novel alkaloid natural products quinazolinone A and B. Based on the structures, the biosynthesis of the compounds most likely involves the conversion of anthranilic acid to anthranilamide, which is subsequently attached to two glycerol units, to produce compounds **1** and **2**. Identification of the biosynthetic gene cluster and subsequent analysis of the individual enzymatic reactions should reveal the precise biosynthetic pathway for these exciting novel molecules.

