## Supplemental Information for "Discovery of Novel Glycerolated Quinazolinones from *Streptomyces* sp. MBT27"

**Figures list**

**Fig. S1** Effect of carbon source utilization on secondary metabolite profiles of *Streptomyces* sp. MBT27

**Fig. S2** Changes in the secondary metabolite profiles of *Streptomyces* sp. MBT27 depending on the glycerol concentration

**Fig. S3** $^1$H NMR spectrum of *quinazolinone A* (**1**) in $CD_3OD$

**Fig. S4** APT spectrum of *quinazolinone A* (**1**) in $CD_3OD$

**Fig. S5** $^1$H-$^1$H COSY spectrum of *quinazolinone A* (**1**) in $CD_3OD$

**Fig. S6** HSQC spectrum of *quinazolinone A* (**1**) in $CD_3OD$

**Fig. S7** HMBC spectrum of *quinazolinone A* (**1**) in $CD_3OD$

**Fig. S8** $^1$H-$^1$H NOSEY spectrum of *quinazolinone A* (**1**) in $CD_3OD$

**Fig. S9** HRMS spectrum of *quinazolinone A* (**1**)

**Fig. S10** UV spectrum of *quinazolinone A* (**1**)

**Fig. S11** IR spectrum of *quinazolinone A* (**1**).

**Fig. S12** $^1$H NMR spectrum of *quinazolinone B* (**2**) in $CD_3OD$

**Fig. S13** APT spectrum of *quinazolinone B* (**2**) in $CD_3OD$

**Fig. S14** $^1$H-$^1$H COSY spectrum of *quinazolinone B* (**2**) in $CD_3OD$

**Fig. S15** HSQC spectrum of *quinazolinone B* (**2**) in $CD_3OD$

**Fig. S16** HMBC spectrum of *quinazolinone B* (**2**) in $CD_3OD$

**Fig. S17** $^1$H-$^1$H NOSEY spectrum of *quinazolinone B*(**2**) in $CD_3OD$.

**Fig. S18** HRMS spectrum of *quinazolinone B* (**2**)

**Fig. S19** UV spectrum of *quinazolinone B* (**2**)

**Fig. S20** IR spectrum of *quinazolinone B* (**2**)

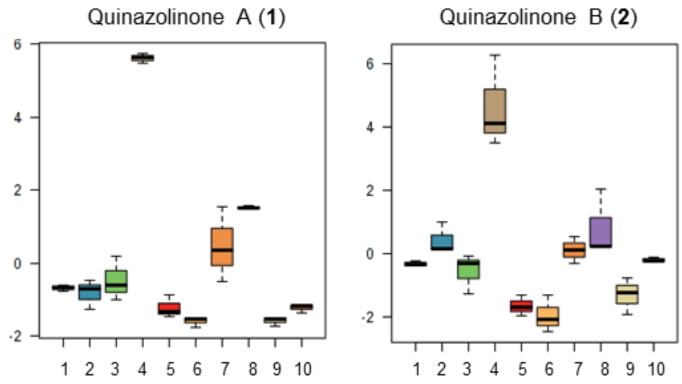


**Fig. S1 Effect of carbon source utilization on the secondary metabolite profiles of *Streptomyces* sp. MBT27.** Box plots show the relative intensities of quinazolinones A (**1**) and B (**2**) in cultures grown in MM with different carbon sources, namely: 1. mannitol 1% + glycerol 1%; 2. GlcNAc 1%; 3. glycerol 1%; 4. glycerol 2%; 5. mannitol 1%; 6. mannitol 2%; 7. glucose 1%; 8. glucose 2%; 9. arabinose 1%; 10. fructose 1%. Note the spectacular increase in production when the glycerol concentration is doubled.

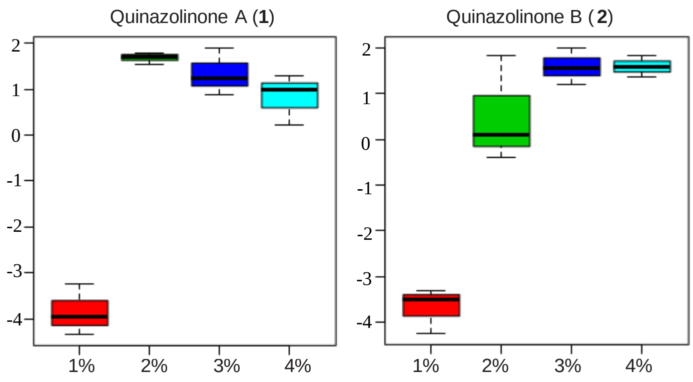


**Fig. S2** Changes in the secondary metabolite profiles of *Streptomyces* sp. MBT27 depending on the glycerol concentration. Box plots show the relative intensities of quinazolinones A (**1**) and B (**2**) in the cultures with added 1%, 2%, 3% or 4% (w/v) glycerol.

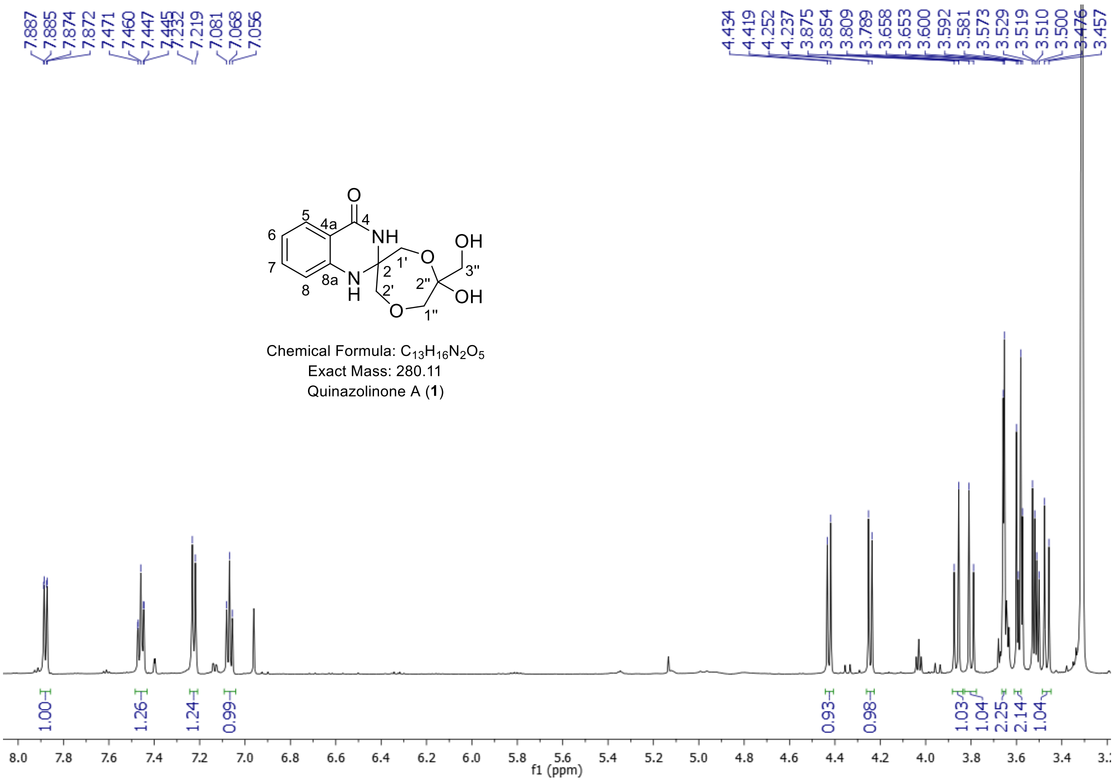


**Fig. S3** $^1$H NMR spectrum of *quinazolinone A* (**1**) in $CD_3OD$

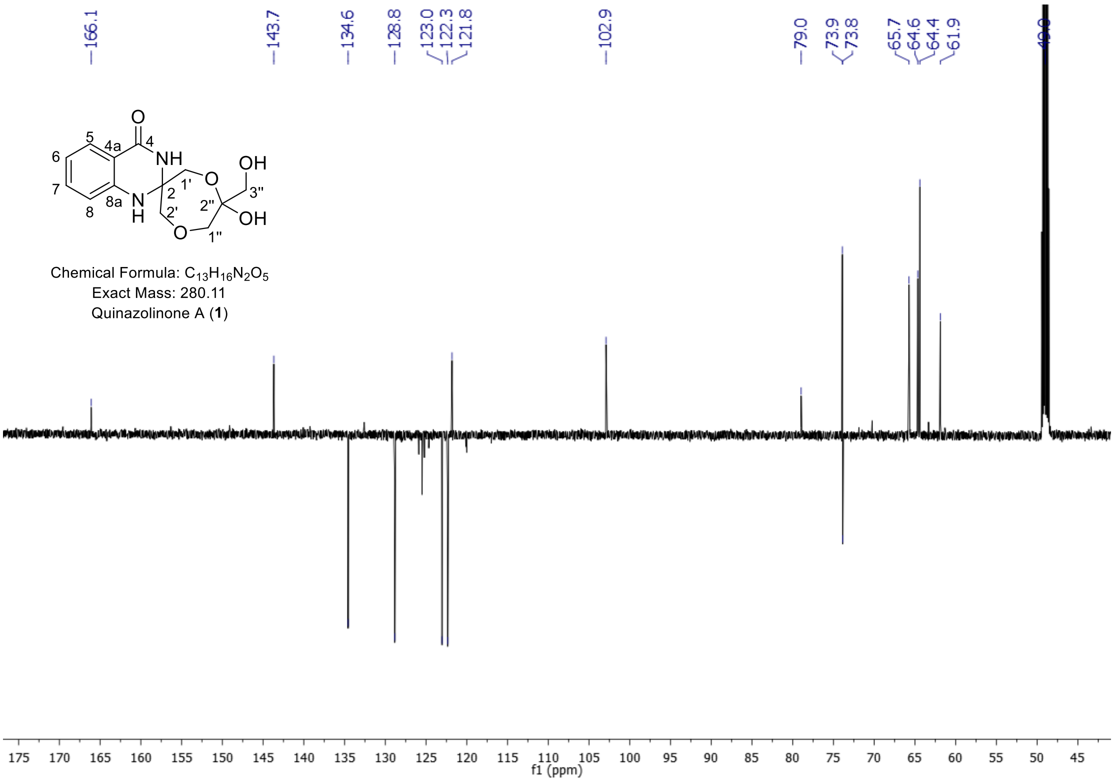


**Fig. S4** APT spectrum of *quinazolinone A* (**1**) in $CD_3OD$

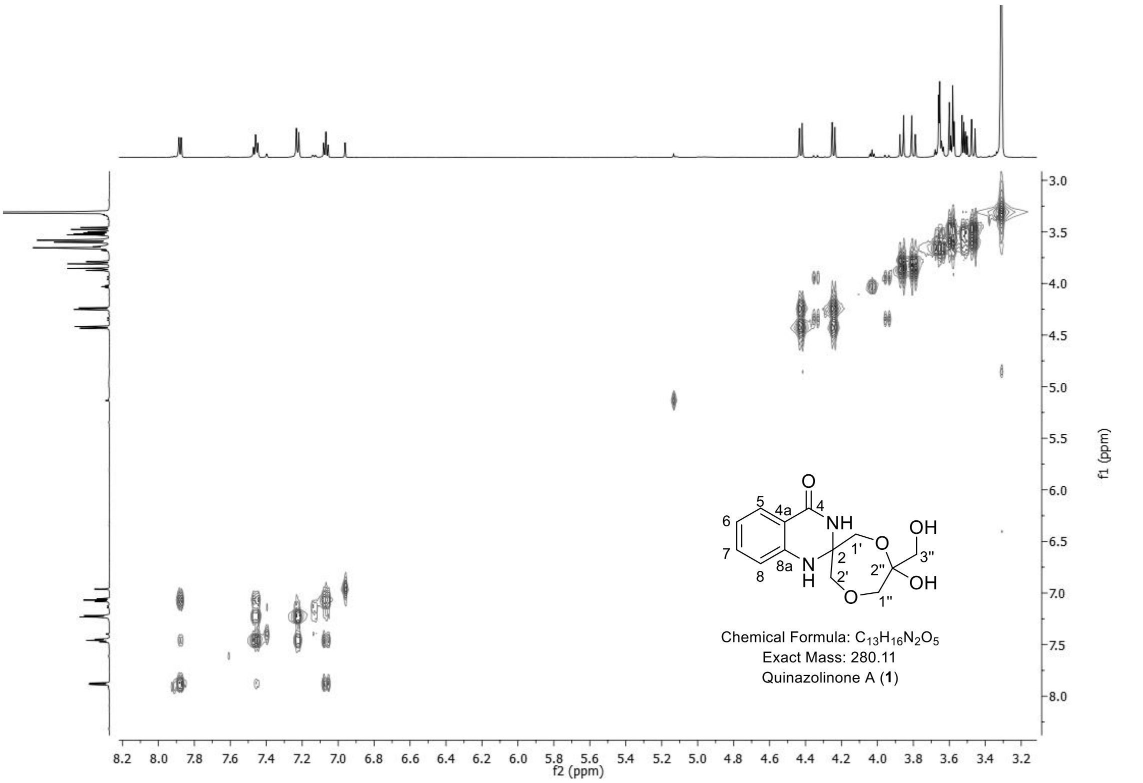


**Fig. S5** $^{1}$H-$^{1}$H COSY spectrum of *quinazolinone A* (**1**) in $CD_3OD$

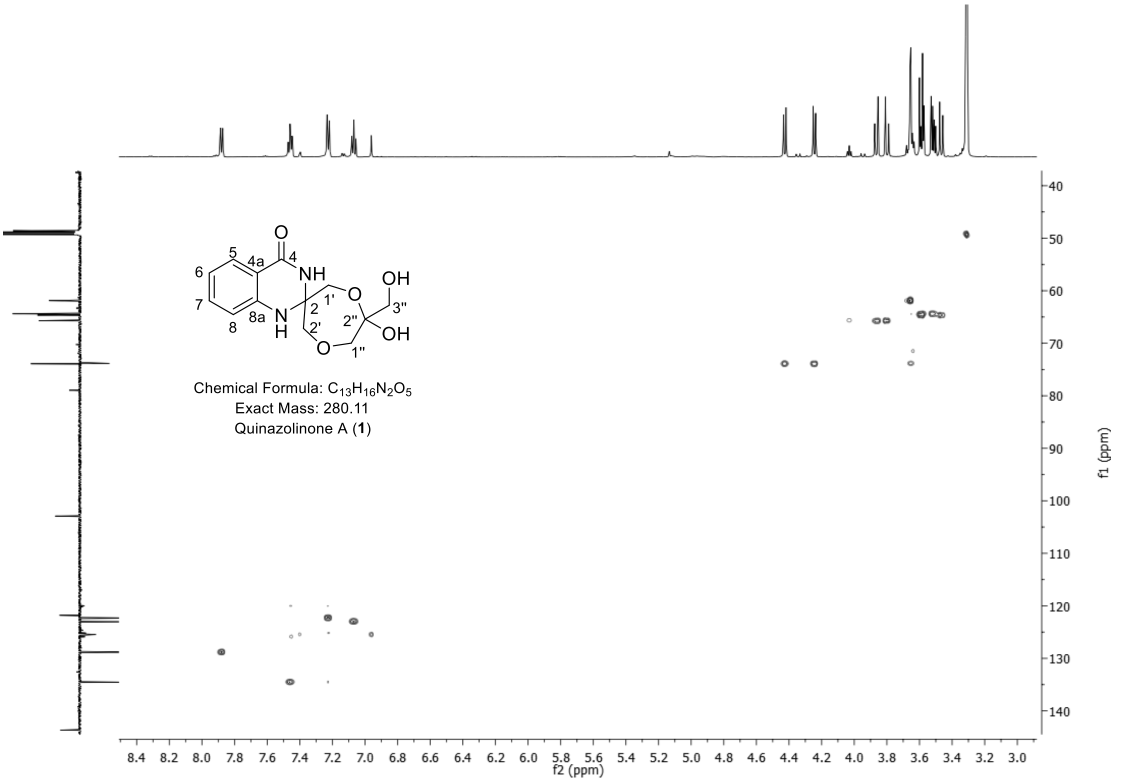


**Fig. S6** HSQC spectrum of *quinazolinone A* (**1**) in $CD_3OD$

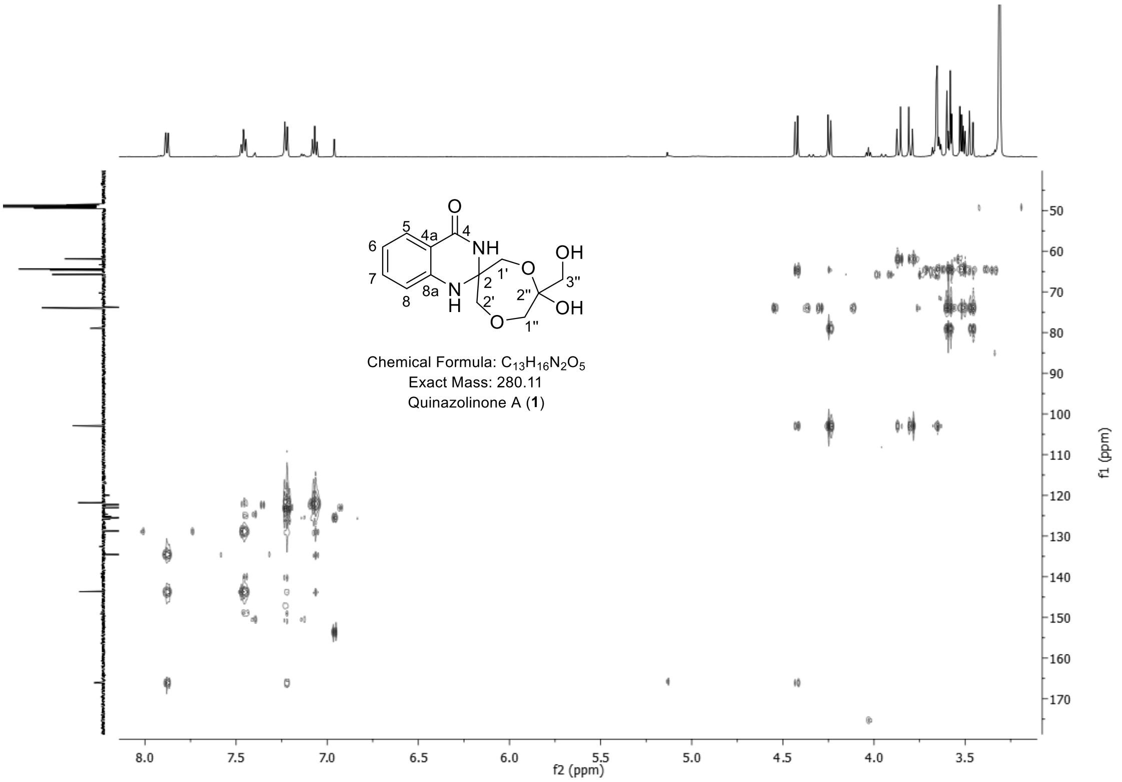


**Fig. S7** HMBC spectrum of *quinazolinone A* (**1**) in $CD_3OD$

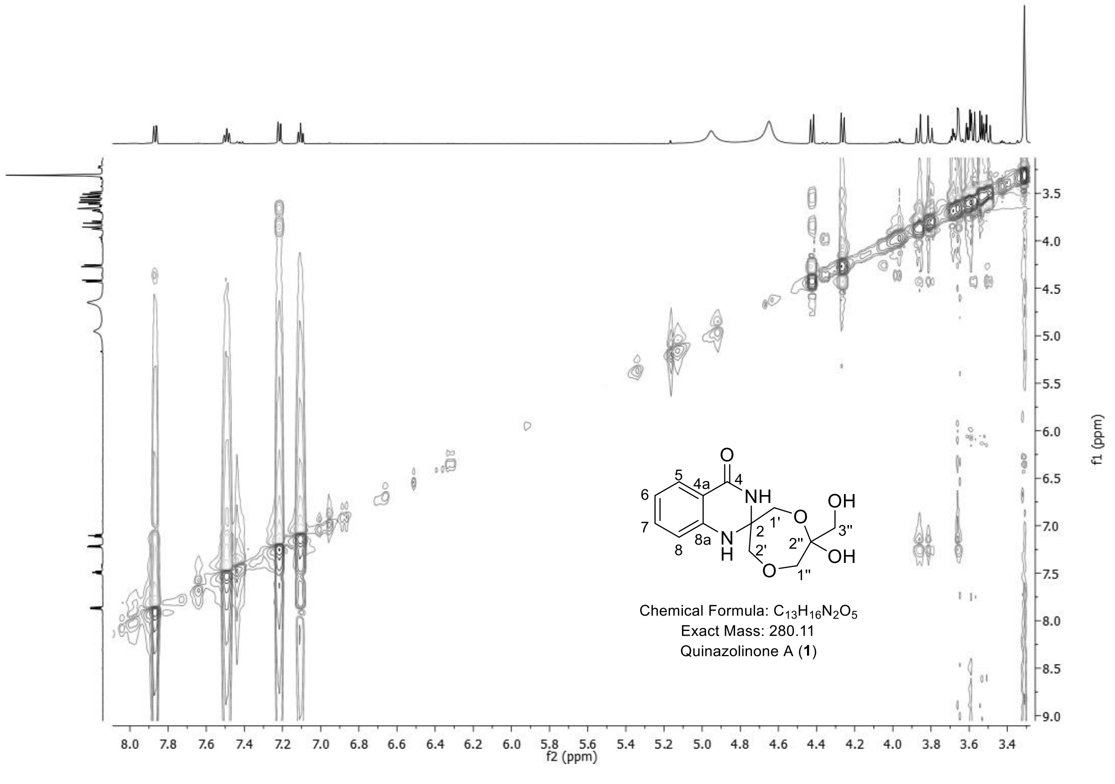


**Fig. S8** $^1H$-$^1H$ NOSEY spectrum of *quinazolinone A* (**1**) in $CD_3OD$

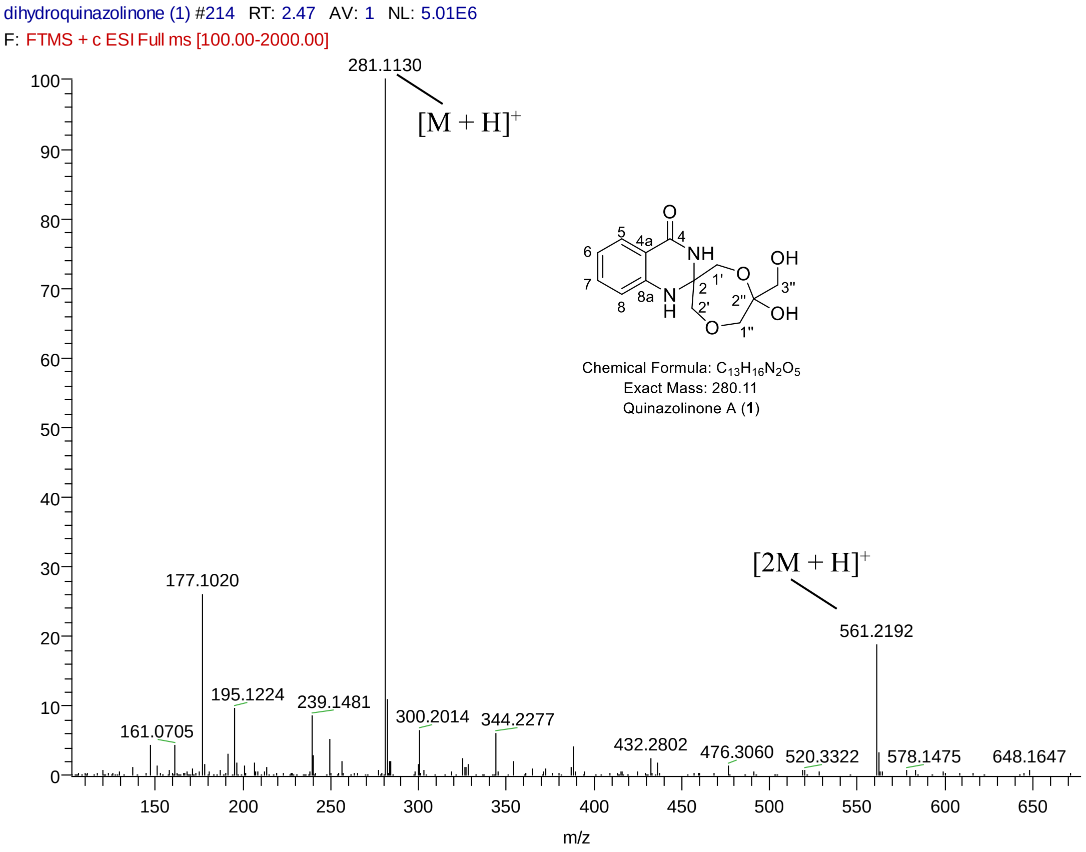


**Fig. S9** HRMS spectrum of *quinazolinone A* (**1**)

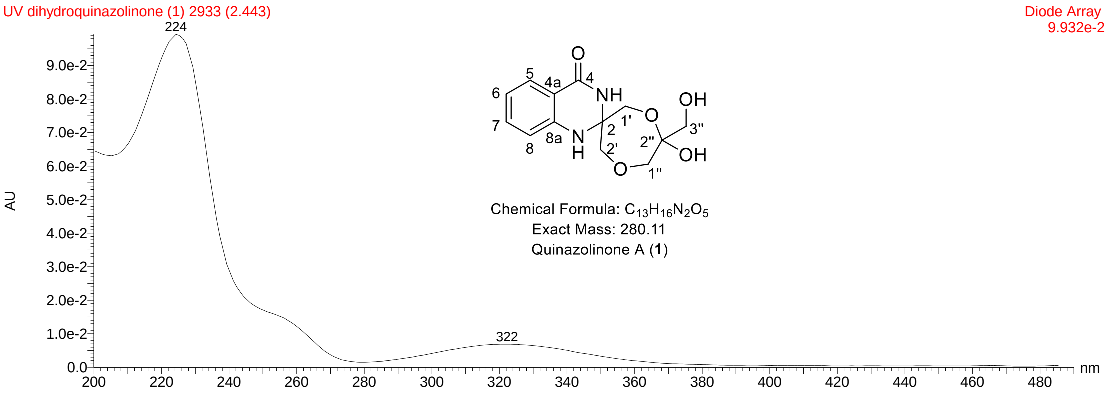


**Fig. S10** UV spectrum of *quinazolinone A* (**1**)

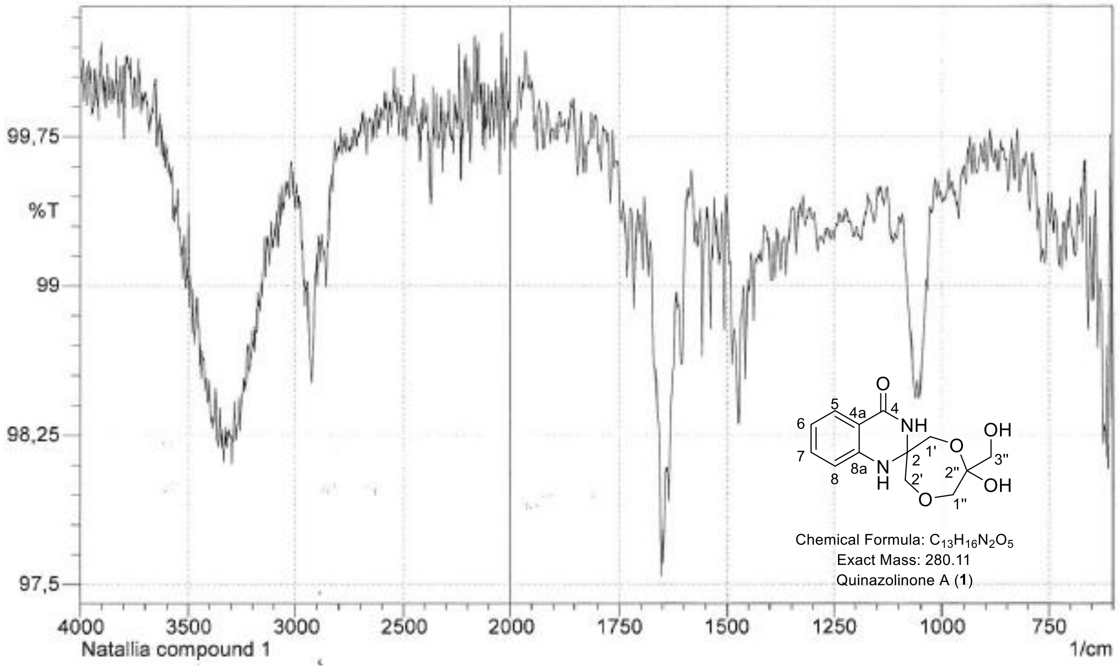


**Fig. S11** IR spectrum of *quinazolinone A* (**1**)

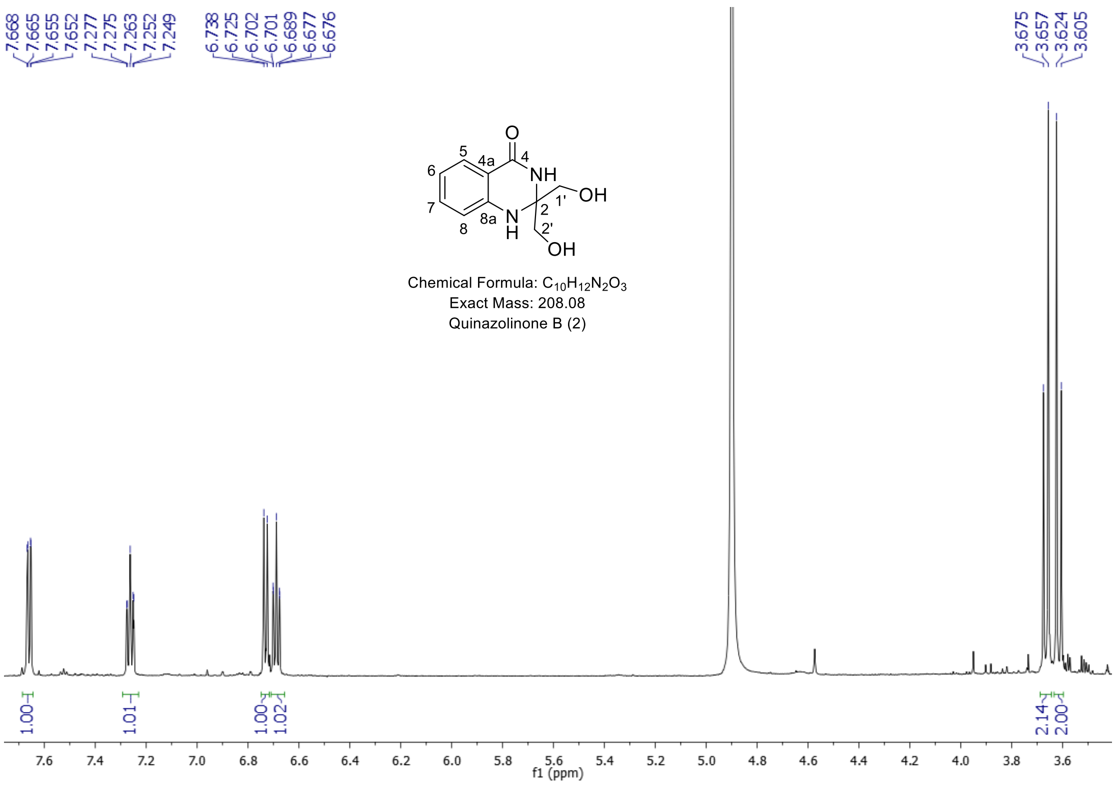


**Fig. S12** [1]H NMR spectrum of *quinazolinone B* (**2**) in $CD_3OD$

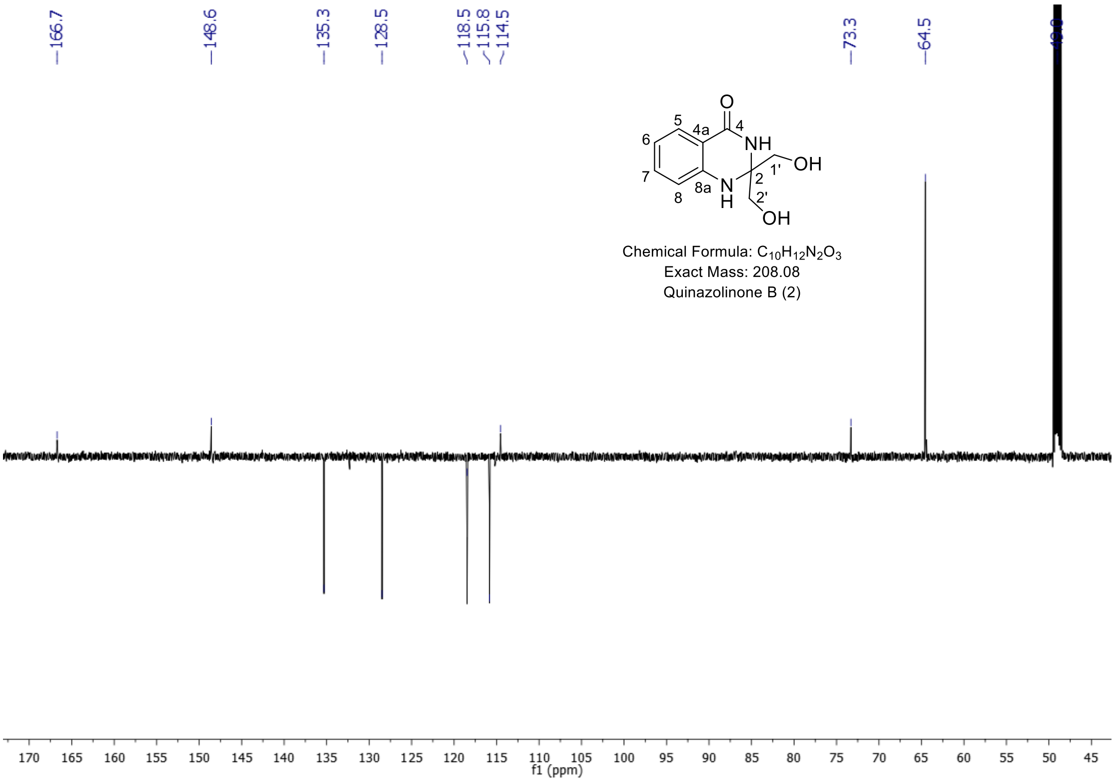


**Fig. S13** APT spectrum of *quinazolinone B* (**2**) in $CD_3OD$

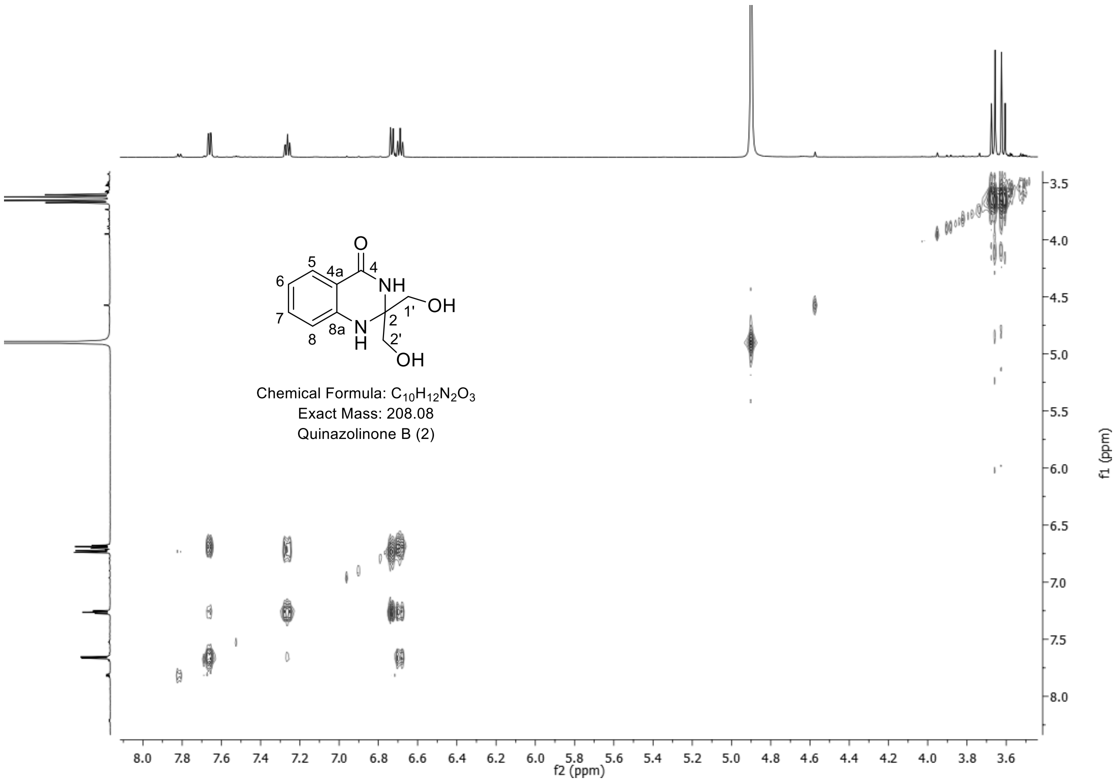


**Fig. S14** $^{1}$H-$^{1}$H COSY spectrum of *quinazolinone B* (**2**) in $CD_3OD$

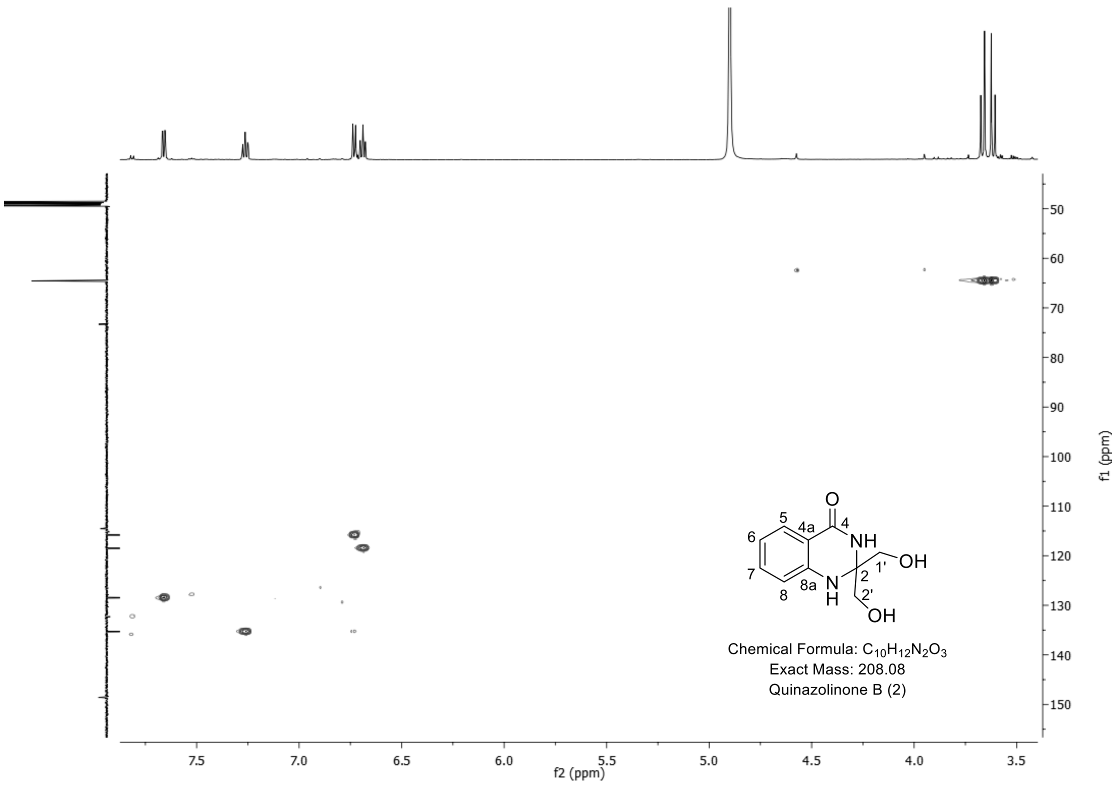


**Fig. S15** HSQC spectrum of *quinazolinone B* (**2**) in $CD_3OD$

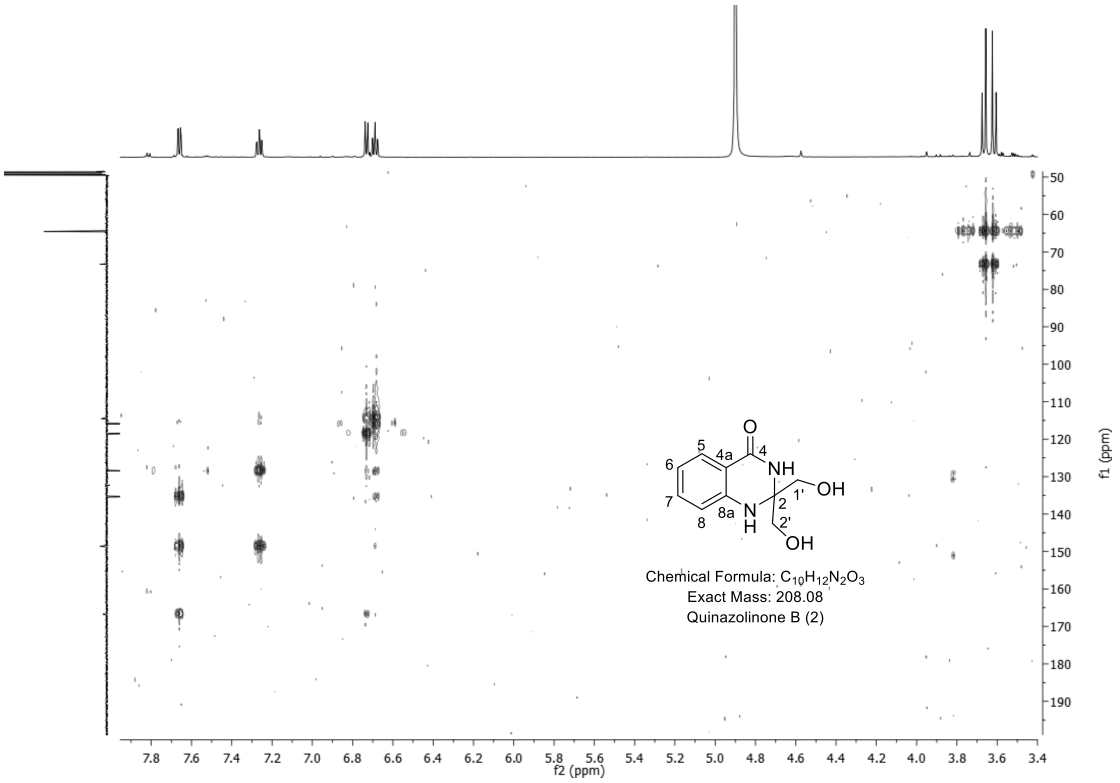


**Fig. S16** HMBC spectrum of *quinazolinone B* (**2**) in $CD_3OD$

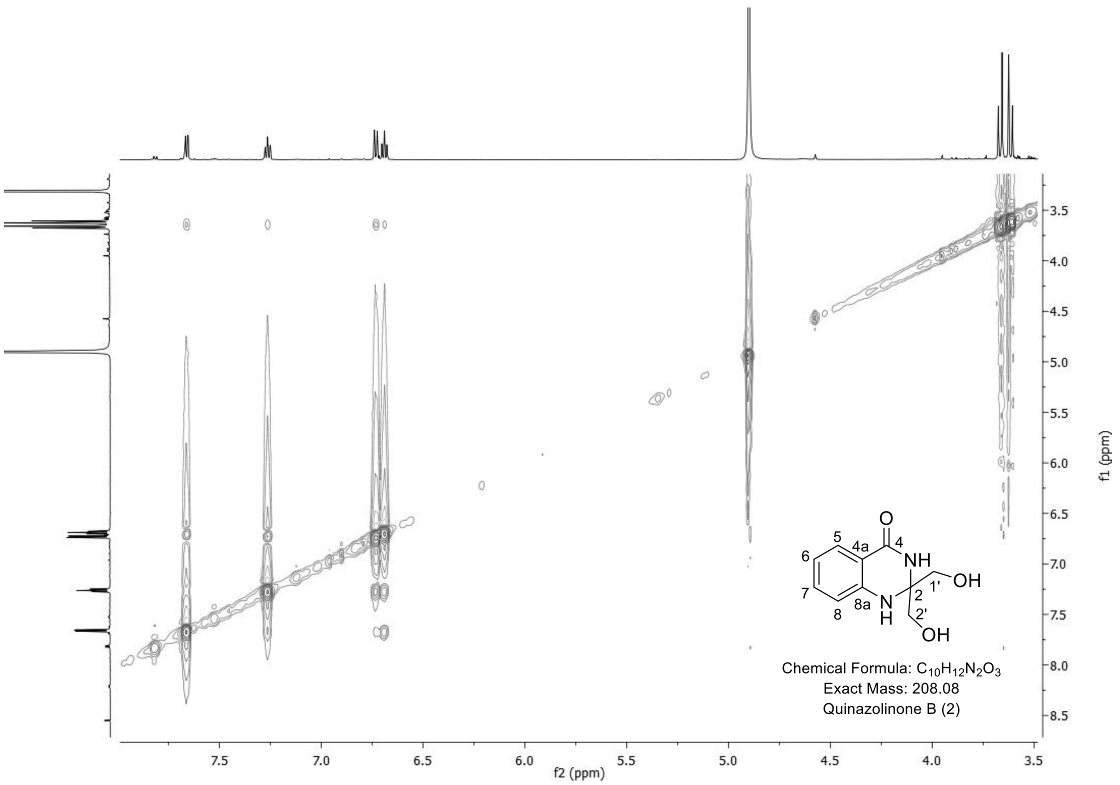


**Fig. S17** [1]H-[1]H NOSEY spectrum of *quinazolinone B* (**2**) in $CD_3OD$

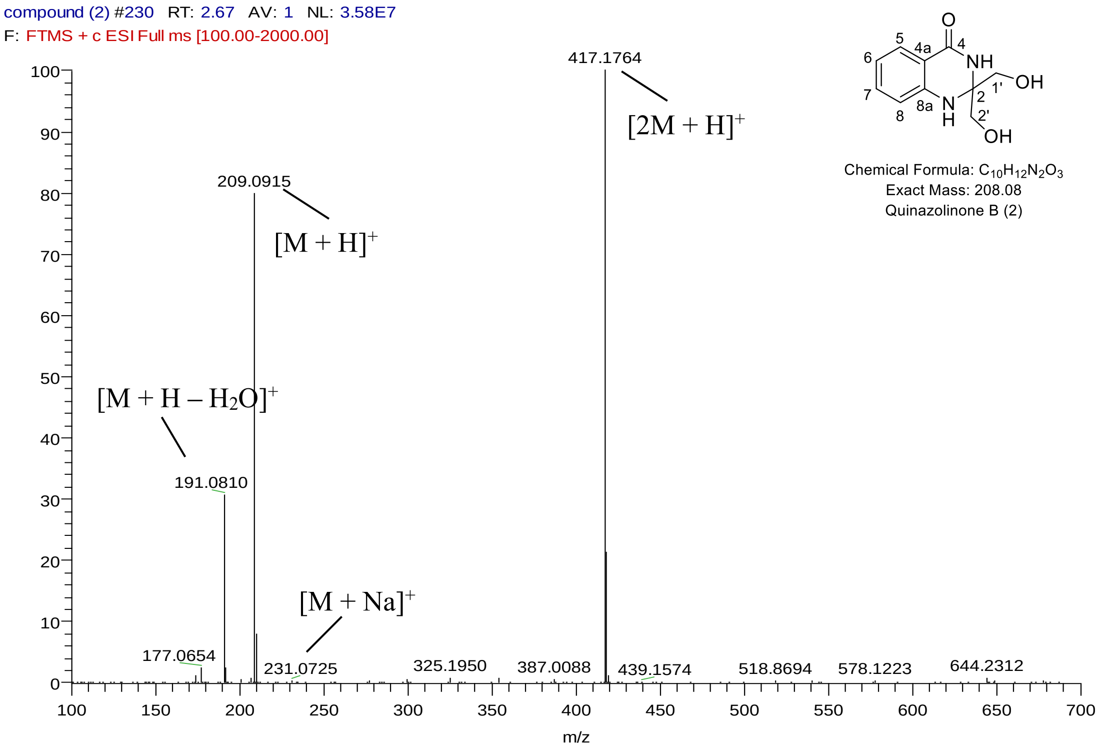


**Fig. S18** HRMS spectrum of *quinazolinone B* (**2**)

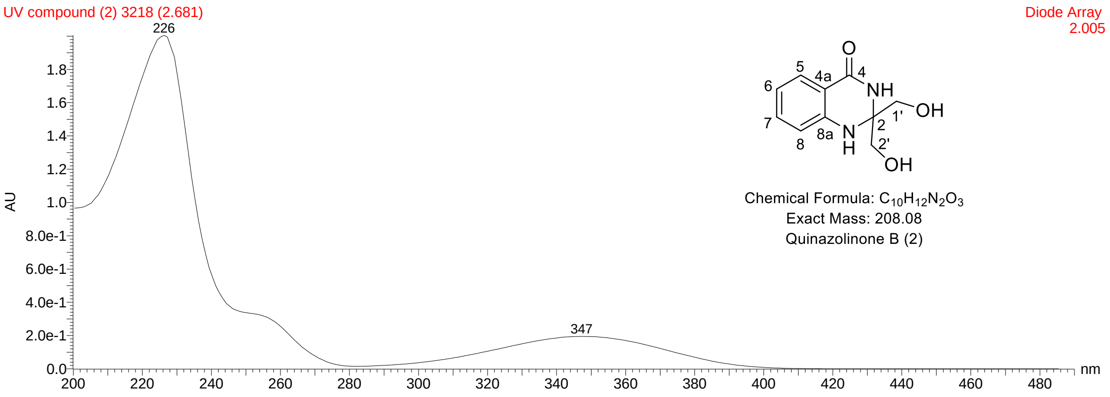


**Fig. S19** UV spectrum of *quinazolinone B* (**2**)

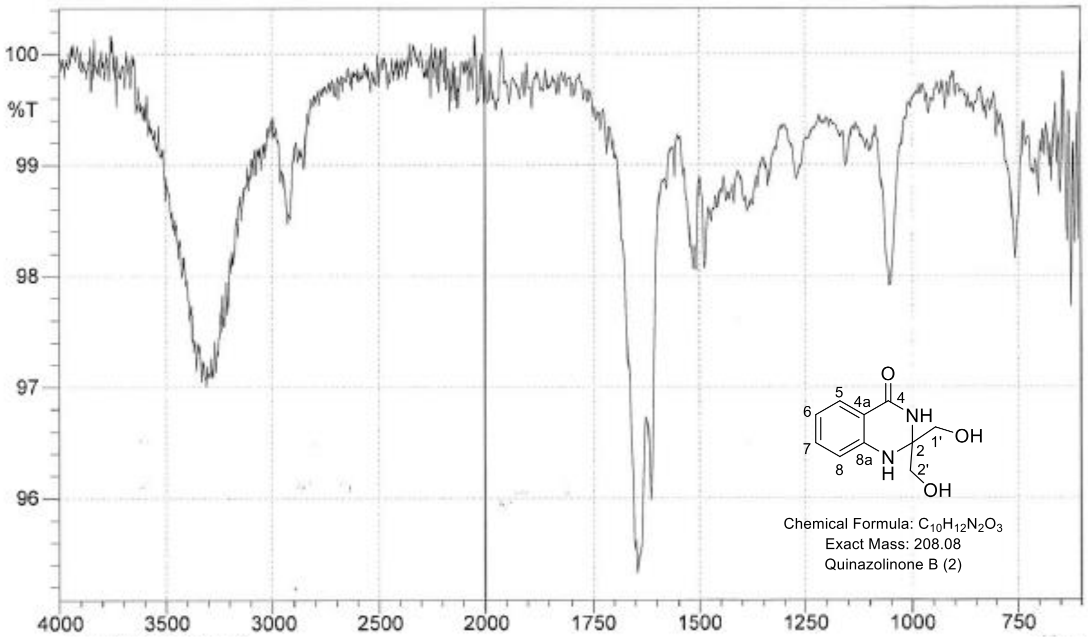


**Fig. S20** IR spectrum of *quinazolinone B* (**2**)
